# CoFrEE: An Application to Estimate DNA Copy Number from Genome-wide RNA Expression Data

**DOI:** 10.1101/2023.08.25.554898

**Authors:** Anita Gaenko, Dipankar Ray, Derek J. Nancarrow

## Abstract

We introduce CoFrEE, a simple python-based approach to extracting copy number data from expression values that works with either RNAseq or array-based expression data. CoFrEE works best in tumor cohorts that include a subset of non-tumor tissues and is applied to processed (RSEM, RPKM or TPM) expression, rather than raw data. Experiments with real public data suggest CoFrEE can provide copy number estimations comparable to existing RNAseq-based approaches, with the advantage of also being applicable to the multitude of older expression-array cohorts.

## INTRODUCTION

RNA-based data make up the bulk of archival cancer-related cohorts, for example in the Gene Expression Ombudsmen (GEO), yet DNA copy number (CN) alterations involving the loss or gain of multiple consecutive genes, are the key hallmark for most somatic cancer types [1]. Cancer driver genes, often classified as either tumor suppressors genes (TSGs: frequently lost) or oncogenes (frequently gain copies), represent the targets of these CN changes, however, pan-cancer studies show that the expression levels of most genes reflect regional copy number change [2, 3]. Our goal was to leverage the large number of samples stored in historical mRNA cohorts to investigate cancer type specific CN changes for patterns of association between subsets of TSGs and oncogenes. These patterns of broad chromosomal change will act as signposts for key cancer biology subtypes, but characterization will require large sample numbers. While several methods exist to identify CN changes in DNA based (CGH array or WES/WGS) data, we and others (CNVkit-RNA: [4], SuperFreq: [5], CaSPeR: [6], CNAPE: [7]) are working to enlist regional changes in gene expression to identify sample-specific differences in copy number to leverage the abundant mRNA-based whole-genome expression data.

<u>Co</u>py number <u>fr</u>om <u>E</u>xpression <u>E</u>stimation (CoFrEE) is unique in providing an intuitively simple approach appropriate for both RNAseq and array-based expression cohorts. To our knowledge, this is also the first such application to focus on facilitating copy number estimates from publicly available processed gene expression-array data; raw files are not required. The core methodology shares recursive median filtering with CaSpER [6] but employs dedicated by-gene pre-processing and by-sample post-processing to achieve final copy number estimates. The preprocessing step shares similarity to CNV-Kit [4], requiring the presence of non-cancer samples (frequently included in such cohorts). The current version targets in GEO, The Cancer Genome Association (TCGA) formats and local user cohorts but could be extended. We include example analyses of array and RNAseq-based cancer cohorts, including comparisons to CGH-based DNA analyses and to CaSpER, confirming utility against existing methodologies.

## FEATURES

CoFrEE uses a linear Python-based Jupyter Labs notebook approach (**Supplemental Figure 1**), allowing users with minimal coding experience to customize notes, code scripts and data visualization to suit their workflow. The notebook template is divided into sections focused on data formatting, pre-analysis normalization, median filtering, post-filtering adjustment (critical for accuracy) and data visualization/export. Functions allows variable median window sizing (gene counts) and iteration count options, as well as the ability to tailor per-genome CN normalization. Example usage of external data visualizations are implemented in template notebooks.

### Gene-based pre-normalization of gene expression

Methodology requires the presence of a normal expression profile, to act as the gene-based denominator for copy-number estimates. Given the tissue-specific nature of gene expression, these data are best provided as a per gene average from accompanying non-cancer samples of the same or similar origins (organ or tissue type) as the cancer cohort. Source data from normalized expression-array formats (e.g. Illumina or Affymetrics genome-wide array studies) or those of RNAseq studies (RSEM, RPKM or TPM) provide input. Since the first step of the analyses is to generate a per-gene log 2 ratio, the specific expression format needs to be consistent and batch normalized with both control and target samples.

Current implementation uses gene expression averaging to generate the ‘reference’ sample, though median or trimmed mean approaches may also be appropriate.

### Recursive median filtering of gene-based expression values

The Scipy (version 1.6.3) function *signal*.*medfilt* was implemented for median filtering and empirical testing determined that three layers of median filtering was optimal. As a feature of this algorithm, odd-numbered window sizes are preferred, centering distribution around each datapoint. We suggest an increasing incremental median-filter window-size (e.g. 11, 21 and 41), however, static window sizing (e.g. 9, 9, 9) can be used. The ability to export each filtered layer provides opportunity for users to test and refine window size and median filtering layer depth as needed. A variance-based median smoothed algorithm may improve CN data quality but would need application-specific (expression platform) testing.

### Genome-based post-normalization and cytoband CN determination

Resulting median-filtered copy number estimations require per-sample normalization to adjust for a) technical factors such as RNA loading, linear amplification variability and normalization quirks, as well as b) genome-wide sampling factors related to sample ploidy and cell-type subpopulations (ploidy heterogeneity). Tetra (4n) or higher base-ploidy can be a feature of cancer genomes, which can skew both DNA and RNA based aneuploidy estimates. CoFrEE offers two systems to adjust for this: average X-chromosome (default), and average autosomal signal. The first is based on evidence that diploid-based mammalian genomes specifically control X-linked gene expression via X-inactivation transcription control, regulating 85% of X-chromosome genes [8], such that affected genes are restricted to a single (mosaic) expressing copy per 2n autosome compliment [9, 10]. Extensive X-chromosome losses are rare for most common cancer-types, and average X-chromosome adjustment provides a gender-neutral, per-genome adjustment to CN-estimates. We provide the alternative average autosome method for instances where an X-inactivation-based approach may not be appropriate, for example, when sex chromosome data was not included in the public release.

We have used cytoband-averaging as our main comparison approach since it emphasizes the approximate nature of CN-estimations, provides an historically relevant framework (cytogenetics) for cross-cohort comparison, and maximizes the fraction of bands (∼ genome coverage) retained from lower resolution array-based platforms. The presented cytobands are based on single band patterns visible on metaphase chromosomes and do not include the sub-band resolutions visible at earlier stages of mitosis [11]. This approach could be adjusted by substituting a user-preferred genome-boundary banding file, for example, a system based on standardize segment-length.

### Data visualization and tabulation features

CoFrEE outputs as flat CSV files containing cytoband averaged and individual gene-level values that approximate log2 adjusted relative copy number. These data can be visualized in different formats and we provide example outputs linked to the genome-wide Tagore cytoband visualization (https://pypi.org/project/tagore/) (**Figure 2A**), Seaborn heatmaps for chromosome-level comparisons (**Supplemental Figure 2B**), and custom UCSC browser tracks for each individual samples (**Supplemental Figure 2C**). **Supplemental Figure 2C** also demonstrates how additional custom data can be added via additional UCSC browser data tracks to enhance key locus identification. We included UCSC tracks for cancer genes extracted from Cancer Gene Census (CGC) [12] and DORGE [13] datafiles, which respectively apply experimentally determined and predictive approaches to oncogene/tumor suppressor identification.

### Comparison analyses

We provide two example studies as comparisons. In **Figure 1A** we use Tagore cytoband visualizations library (https://pypi.org/project/tagore/) to contrast CGH-based DNA copy number with CoFrEE estimated copy number in the esophageal adenocarcinoma (EAC; n=88) cancer samples within the ESCA TCGA dataset [14]. **Figure 1A** demonstrates the general consistency of CGH-DNA and CoFrEE-based CN data with historically consistent regions of CN losses (chromosomes 4q, 5q, 9p, 16q, 17p, 18q and 21q) or gains (7p, 7q, 8q, and chromosome 20) in EACs. Cytoband changes can be seen in several genomic regions, including chromosome 13q, unique to CoFrEE, and focal regions on 16p and 18p unique to the DNA CGH analysis. **Supplemental Figure 3** provides a semi-quantitative summary comparison between ‘real’ CGH-based and CoFrEE estimated copy number for both EAC and ESCC esophageal cancer types, indicating strong categorical agreement between real and mRNA-estimated copy number (loss, gain or neutral assignments) for greater than 90% of cytobands copy number averages, but provided detailed analyses do show that CoFrEE has a tenancy to over-report large CN change and miss narrow regions of change, such as the gain at centromeric 18p focused on GATA6 (ref).

**Figure 1:**
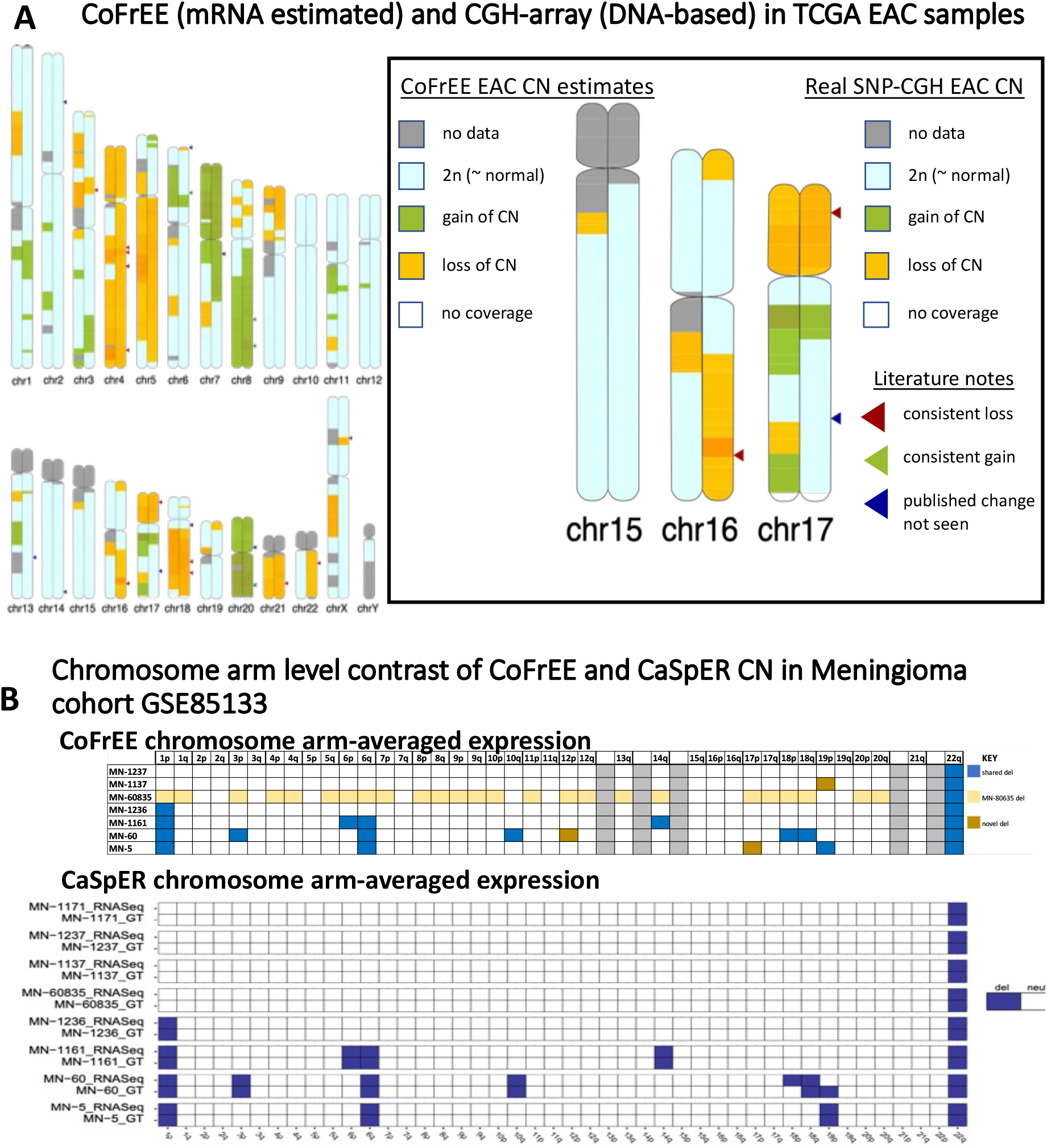
A) Comparison of real DNA-based CGH and mRNA-base CoFrEE estimates of copy number in EAC sample (n=88) from the TCGA ESNA cohort of esophageal cancers. We used Broad Firehose, GISTIC generated CGH copy number data, and CoFrEE analysis using per-gene averaged expression RSEM values from 11 non-cancer squamous esophageal samples from cancer patients as our control reference group, followed by tree median filtering layers of 11, 21, and 41 as described. We then used X-chromosome averaged per-sample adjustment. For both CGA and CoFrEE estimates we generated per-cytoband averages by determining mean per-cytoband gene levels per sample, then averaging EAC samples. For the CGH copy number estimates GISTIC thresholds of 0.2 were used to designate gain (>0.2) or loss (<-0.2) from regions of 2n (between 0.2 and -0.2) log2 ratio units, Since CoFrEE has a more sensitive baseline (as described in the text) we used our standard gain/loss thresholds of 0.25 and -0.25 respectively. Grey and clear regions represent cytobands with data for less than 10 or no genes by real CN or CoFrEE, respectively. B) Comparison of mRNA based CoFrEE and CaSpER copy numbers in meningioma samples with chromosome 22 deletions. GEO accession GSE85133, which includes both mRNA expression and DNA-based NGS copy number data provided as part of the CaSpER R package [6]. We used the by-gene average of samples without losses (by NGS DNA copy number) on chromosome 22 (containing the key region of loss for meningioma; [15]) as the reference for samples with chromosome 22 deletion, exactly as presented in Harmanci 2020:Fig 3 [6]). We used both B) chromosome arm-averaged and Supplemental Figure 4) genome-wide heatmap-based to demonstrate that CoFrEE (top) similarly identifies regions of sample specific changes to CaSpER (bottom) analyses (indicated with arrows). CoFrEE shows some oversensitivity, highlighted in tan (weak) and brown (strong) areas of deletion not seen by CaSpER.

We show a p or q arm-based comparison for a meningioma cohort (GEO accession: GSE85133 [15]) provided as part of the CaSpER R package [6]. We used both tabular (**Figure 1B**) and heatmap (**Supplemental Figure 4**) comparisons to demonstrate the similarity between CN estimation methods, with consistent disease-critical changes on chromosome 22, though again, CoFrEE has a small tendency towards over-sensitive detection of CN change.

## Supporting information

Supplemental Material - example code

## IMPLEMENTATION

CoFrEE 2021 was developed as a Juypter Labs web-browser based application suitable for Windows, Mac OS X, Linux or Unix-based systems, with a simple linear structure (example notebook included as **Supplemental Material** was converted to PDF using Vertopal web converter; https://www.vertopal.com) designed for interrogation of local datasets alongside those in GEO or TCGA. It requires inclusion of a subset of non-cancer tissue samples essential for gene-based normalization. As a method compatible with both expression array and RNA-seq based studies, CoFrEE does not benefit from the inclusion of BAF data, estimated from RNAseq allelic expression, as used by CaSpER and CNVkit-RNA. Therefore, CoFrEE will generate lower-resolution copy number data and requires each detected CN change to include multiple genes (our minimal threshold was 10 genes). This trade-off comes with the ability to apply CoFrEE to larger number of pre-existing, disease-specific datasets, providing potential for an expression-based meta-analysis of CN changes.

## ACKNOWLEDGEMENTS

We wish to thank Prof Arvind Rao and Visweswaran Ravikumar for helpful discussion.

## Abbreviations

CN: [DNA copy number]
BE: [Barrett’s esophagus]
EAC: [esophageal adenocarcinoma]
ESCC: [esophageal squamous cell carcinoma]

## SUPPLEMENTAL MATERIAL

**Supplemental Figure 1:**
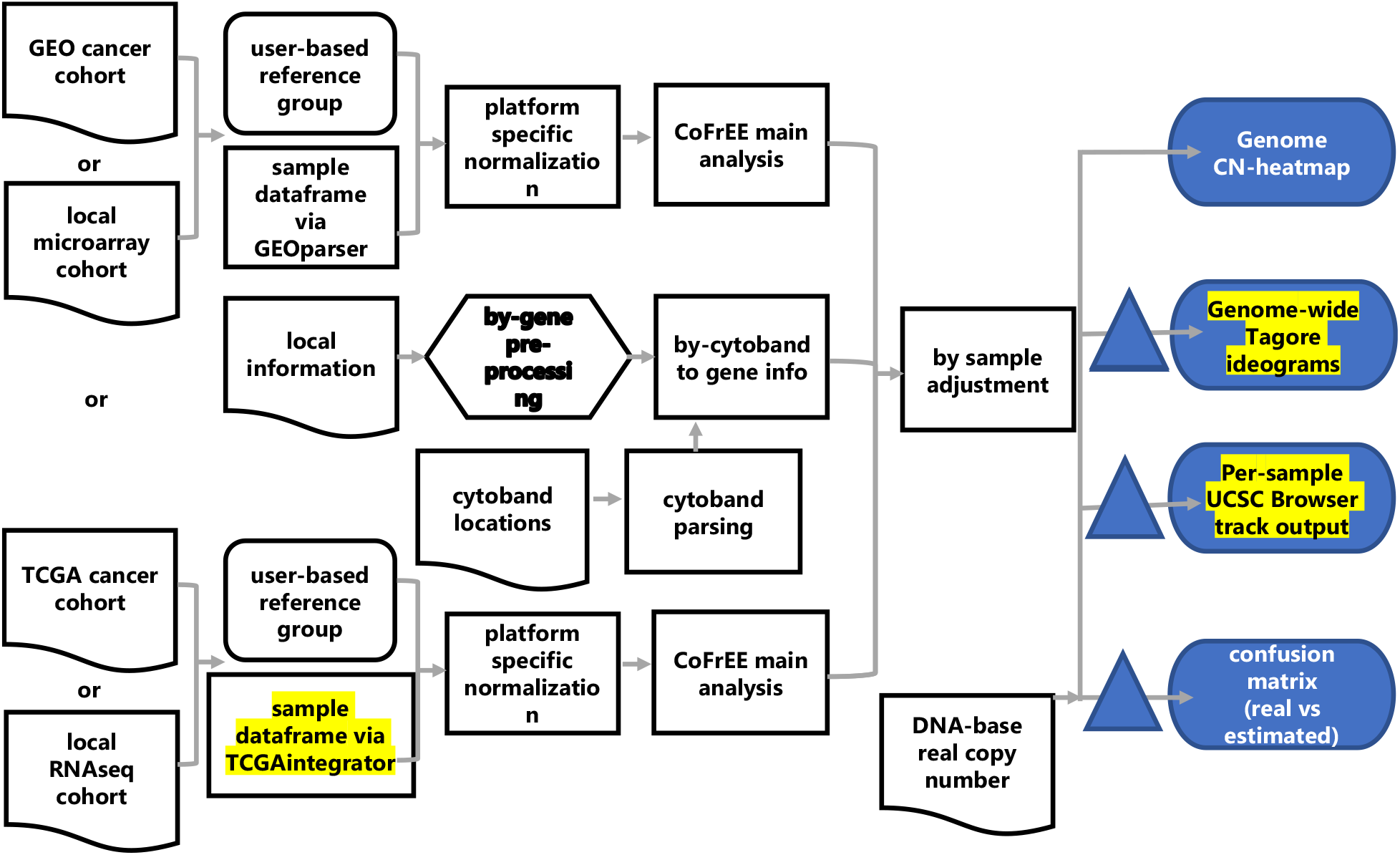
Schema of CoFrEE analysis. GEO, TCGA or similar local datasets in CVS format are read in, along with gene to genome location and cytoband to genome location files. User chosen track is used as a baseline for per-gene normalization and log2 transformation. Recursive median filtering is then used to generate region-smoothed per-gene relative copy number values that are then averaged to cytoband. Resulting CVS files for gene-specific and cytoband-averaged CN estimates can then be input into down-stream applications, with examples of Tagore, Heatmap, UCSC browser and tabulated comparison provided.

**Supplemental Figure 2:**
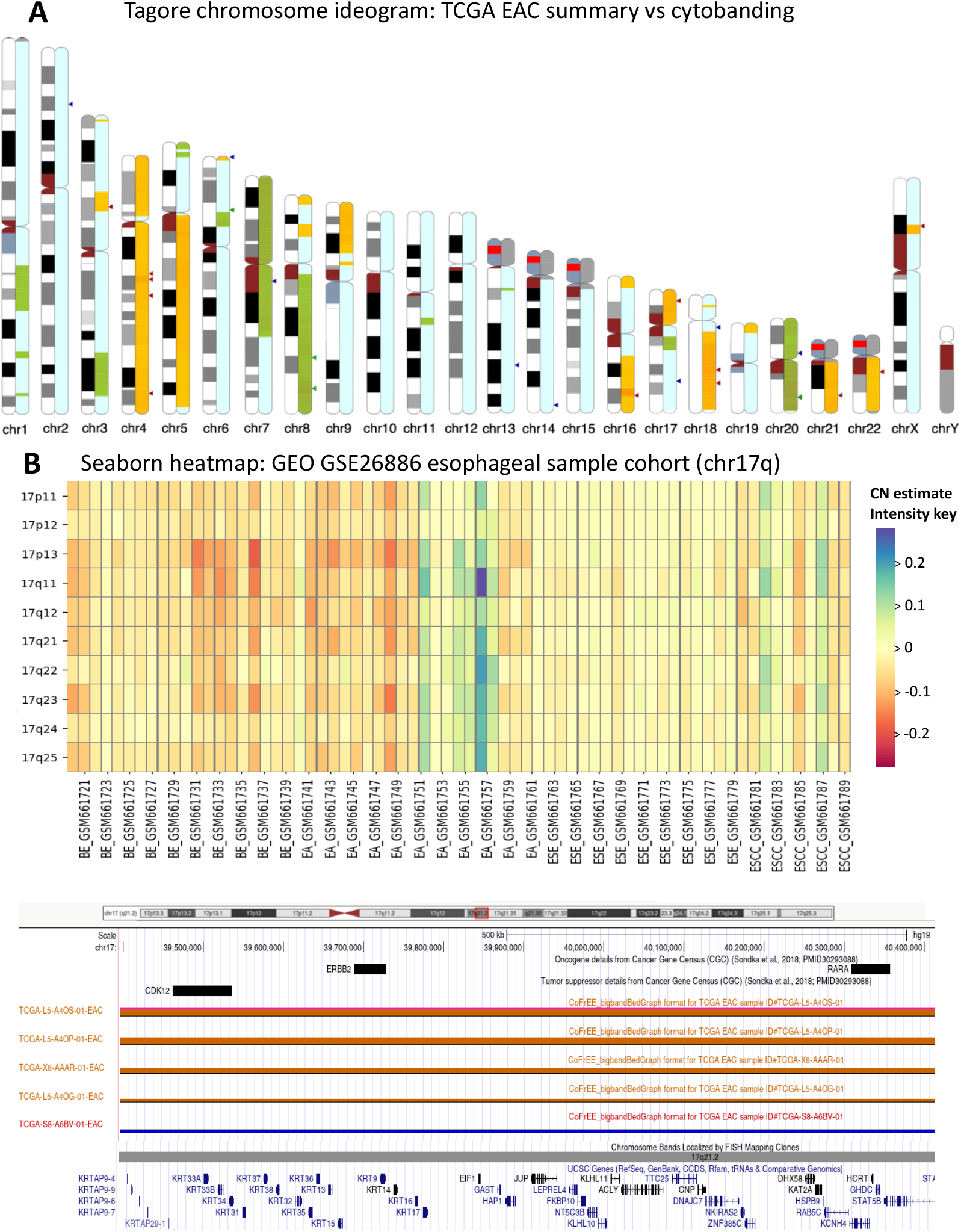
Demonstration visuals from CoFrEE data. **A)** Tagore python library to present genome-wide snapshot showing cytobands (**left**) with copy number estimates (**right**). Cytoband colorations represent best-fit Geimsa (G) banding patterns, while estimated CN colors result from using CoFrEE estimates of < -0.25 (loss), > 0.25 (gain) and between these values as 2n. Cytobands with less than 10 datapoints (genes) are shown in grey (no data) while cytobands with no genes are clear. **B)** Seaborn python library generated heatmap of samples from GEO cohort GSE26886 (Wang et al., 2012), ordered by tissue type and colored by CN estimate. **C)** custom UCSC Browser tracks for five example EAC samples, showing the chromosomal region of 17q21 surrounding ERBB2, an oncogene known to be amplified in a subset of EACs. The feature-rich UCSC Browser format offers a well-known interface with a broad range of functionality, including the ability to easily re-order samples, zoom, and combine additional custom data-tracks. Additional custom tracks shown (top) containing known oncogenes and tumor suppressors from Cancer Gene Census (CGC) database [12] provide signposts for genes of interest.

**Supplemental Figure 3.**
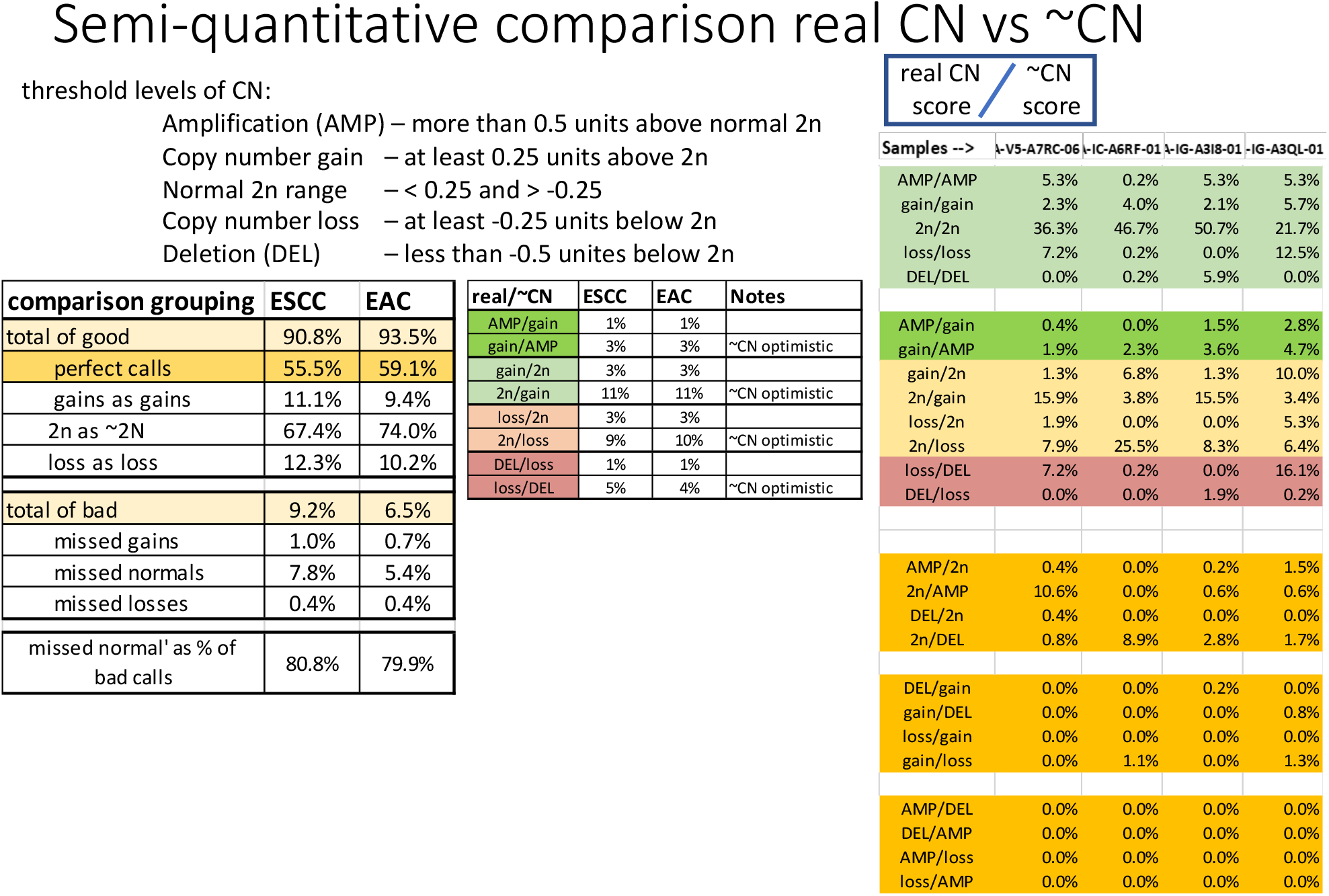
Comparison of real DNA-based CGH and mRNA-base CoFrEE estimates of copy number from the TCGA ESCA cohort of esophageal cancers. Comparison of real DNA-based CGH and mRNA-base CoFrEE estimates of copy number in EAC sample (n=88) from the TCGA ESNA cohort of esophageal cancers. We used Broad Firehose, GISTIC generated CGH copy number data, and CoFrEE analysis using per-gene averaged expression RSEM values from 11 non-cancer squamous esophageal samples from cancer patients as our control reference group, followed by tree median filtering layers of 11, 21, and 41 as described. We then used X-chromosome averaged per-sample adjustment. For both CGA and CoFrEE estimates. We then generated per-cytoband averages of genes per sample, based on UCSC cytoband boundary locations, then averages samples for each cytoband for the final comparison values. For the CGH copy number estimates GISTIC thresholds of 0.2 were used to designate gain (>0.2) or loss (<-0.2) from regions of 2n (between 0.2 and -0.2) log2 ratio units, Since CoFrEE has a more sensitive baseline (as described in the text) we used our standard gain/loss thresholds of 0.25 and -0.25 respectively. Grey and clear regions represent cytobands with data for less than 10 or no genes by real CN or CoFrEE, respectively. **Left panel**: Using the above listed gain and loss thresholds for each method to classify cytobands, with <10 loci included in both analyses, as either gains, losses or 2n regions, by either DNA-CGH (real CN) or CoFrEE (∼CN). The number of bands with the same, or different CN classification by each method were then tabulated and categorized as either consistent or inconsistent. The percentage of tabulated subtype-comparisons are shown. **Right panel**: To further characterize the pattern if inconsistent regions, gains/losses were separated into strong or weak via each of the two methods (using 0.5 and -0.5 log2 relative CN units, respectively). The differential comparison patterns where then tabulated, demonstrating that CoFrEE as a tendency towards over-sensitivity, relative to DNA-CGH.

**Supplemental Figure 4.**
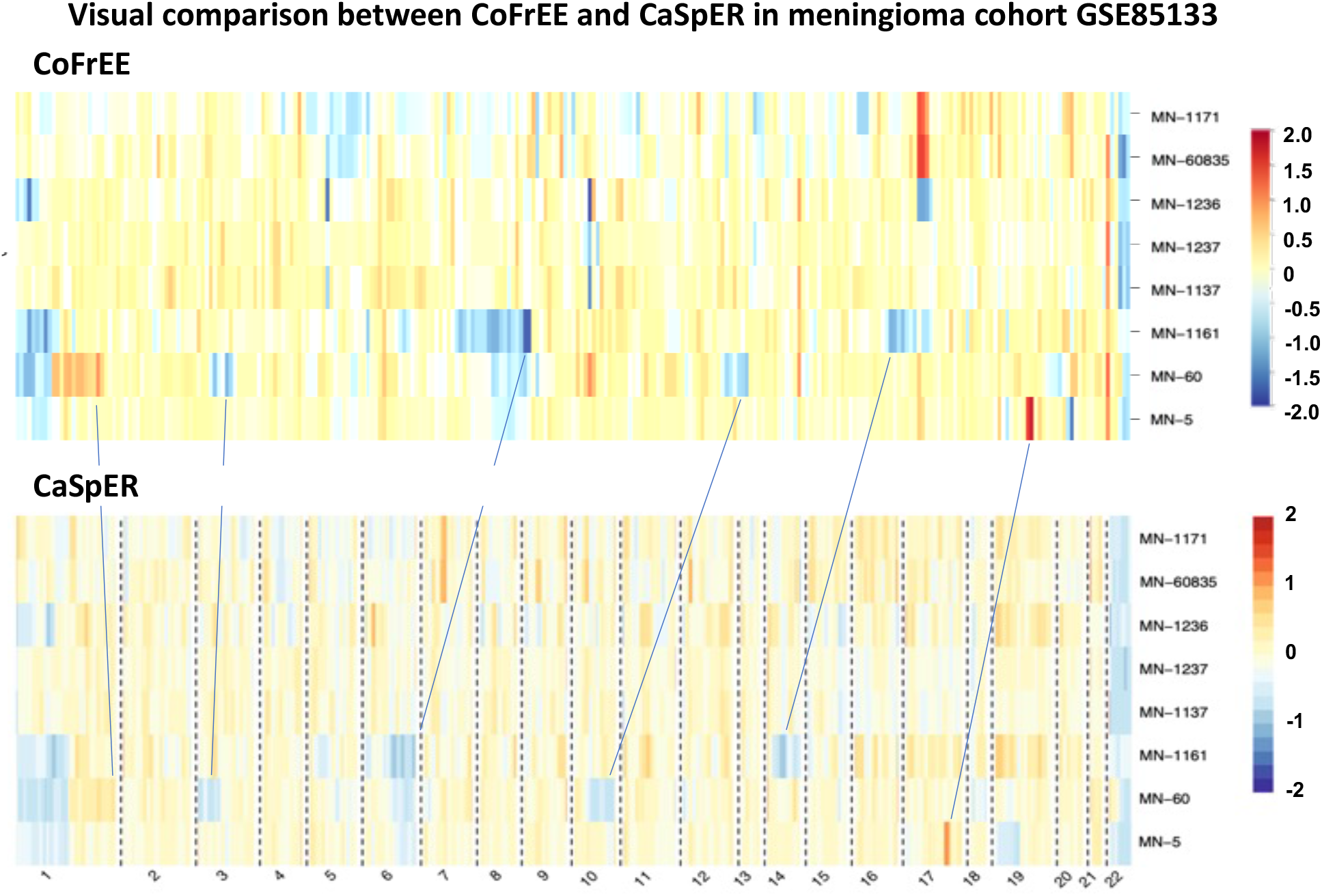
Comparison of mRNA based CoFrEE and CaSpER copy numbers in meningioma samples with chromosome 22 deletions. Comparison of mRNA based CoFrEE and CaSpER copy numbers in meningioma samples with chromosome 22 deletions. GEO accession GSE85133 with both mRNA expression and DNA-based NGS copy number data provided as part of the CaSpER R package (Harmanci 2020) was analyzed. We used the by-gene average of samples without losses (by NGS DNA copy number) on chromosome 22 (containing the key region of loss for meningioma; [15]) as the reference for samples with chromosome 22 deletion, exactly as presented in Harmanci 2020:Fig 3[6]). We show a genome-wide heatmap-based visual as a means to demonstrate that CoFrEE (top) similarly identifies regions of sample specific changes to CaSpER (bottom) analyses (indicated with arrows), although CoFrEE oversensitivity highlights several additional regions in individual samples.

## Notes

Grant support: Supporting grants: RO1CA215596.

Disclosures: The authors report no conflict of interest.

### Competing Interest Statement

The authors have declared no competing interest.

