## Supplemental Material - example code for "CoFrEE: An Application to Estimate DNA Copy Number from Genome-wide RNA Expression Data"

```
# Example CoFrEE code
```

### 1. Initialization

#### 1.1 Global Initializations and Imports

```
import pandas as pd
import numpy as np
from scipy import ndimage, signal, misc
import matplotlib.pyplot as plt
import seaborn as sns

import GEOparse #only needed to access original format example data
```

#### 1.2 Figure Format Controls

This section should probably be moved into section 6 and expanded upon

```
SMALL_SIZE = 8
MEDIUM_SIZE = 10
BIGGER_SIZE = 12

plt.rc('font', size=SMALL_SIZE)          # controls default text sizes
plt.rc('axes', titlesize=SMALL_SIZE)     # fontsize of the axes title
plt.rc('axes', labelsize=MEDIUM_SIZE)    # fontsize of the x and y
labels
plt.rc('xtick', labelsize=SMALL_SIZE)    # fontsize of the tick labels
plt.rc('ytick', labelsize=SMALL_SIZE)    # fontsize of the tick labels
plt.rc('legend', fontsize=SMALL_SIZE)    # legend fontsize
plt.rc('figure', titlesize=BIGGER_SIZE)  # fontsize of the figure
title
```

### 2. Global functions

#### 2.1 cytoband related functions

```
# this can be used if your system allows python to read directly from
a gzipped text file
def getbands(cbfile = "./cytoBand.txt.gz"):
    cband = pd.read_csv(cbfile, names="chr start end cbname
stain".split(), sep="\t")
    cband.cbname = cband.cbname.apply(bigband)
    cband.cbname = cband.chr.apply(lambda x: x[3:]) + cband.cbname
    cbmax = cband.iloc[:,[0,2,3]].groupby(by = "cbname").max()
    cbmin = cband.iloc[:,[1,3]].groupby(by = "cbname").min()
    cband = pd.concat([cbmin,cbmax],axis = 1)
    cband.reset_index(inplace = True)
    return cband
```

```
# cband = getbands("./cytoBand.txt") # original assumed
"cytoBands.txt.gz" was decompressed automatically
'''this loads a standard UCSC Browser cytoband file and reduces
banding pattern
   to a single record per large band with 'start' and 'end' spanning
the combined sub-bands.
   Done to increase the number of cytobands with at least 10 loci, and
as it is consistent
   with the resolution of analysis approach

   columns are 'cbname','chr','start','end','chrnum', consistent with
original
'''
def getbands(cbfile = "./cytoBand.txt"): #manually unpack gz file
    cband = pd.read_csv(cbfile, names="chr start end cbname
stain".split(), sep="\t")
    cband.dropna(inplace=True)
    cband['bigband'] =cband['cbname']
    cband.bigband = cband.bigband.apply(bigband)
    cband.bigband = cband.chr.apply(lambda x: x[3:]) + cband.bigband
#add chr to band designation
    cband['chrnum'] = cband.chr.apply(lambda x: x[3:]) #add ability to
order by chr, then by start
    cband['chrnum'] = cband['chrnum'].replace('X' ,23) #This is human
genome specific!
    cband['chrnum'] = cband['chrnum'].replace('Y' ,24) #This is human
genome specific!
    cband[['chrnum']] = cband[['chrnum']].apply(pd.to_numeric) #This
is dangerous - assumes all strings are numbers!
    cbmax = cband.iloc[:,[0,2,5,6]].groupby(by = "bigband").max() #to
get max location for big bands
    cbmin = cband.iloc[:,[1,5]].groupby(by = "bigband").min() # to get
the min location for bigbands
    cband = pd.concat([cbmin,cbmax],axis = 1) # this destroys the
logical order
    cband.sort_values(by=['chrnum','start'], inplace=True)
    cband.reset_index(inplace = True) # new cband file is ready but
column order needs adjustment
    cband.columns = ['cbname','start','chr','end','chrnum']
    cband = cband[['cbname', 'chr', 'start', 'end', 'chrnum']] #want
to make "Chr" lower case
    cband
    return cband

# bigbands are those visible in late metaphase ... to get these, take
off all the sub-band details after the 'dot'
# example 1p36.3 becomes 1p36
# using bigbands gives a higher density of gene CN reads per band,
providing better unit coverage, at the cost of sensitivity.
def bigband(band):
```

```
    dp = band.find(".")
    if dp == -1:
        return band
    else:
        return band[:dp]

def assign_cytoband(grouped_df, cband, chr_col = "chr", inCHRloc =
"inCHRloc"):
    ''' Receive a dataframe grouped by chromosome
    return the dataframe with cytobands assigned'''
    chrom = f"chr{int(grouped_df.iloc[0][chr_col])}" # convert
chromosome number to name
    grouped_df = grouped_df.copy()
    for cb in cband[cband.chr==chrom].iloc: # for all cytobands in a
given chromosome...
        # what genes have their start inside this cytoband?
        grouped_df.loc[(grouped_df[inCHRloc]>=cb.start) &
(grouped_df[inCHRloc]<cb.end),'cband_start'] = cb.cbname
        grouped_df
    return grouped_df
```

#### 2.2 Pre-smoothing normalization function

```
# pre-smoothing per-gene normalization
# requires users to have chosen and averaged a reference sample
groups, such as non-cancer samples of the same tissue type
# generate per gene ratios and convert to log2 format
def logFC(myFile,ref_col,rel_cols = 7):
    ratios = myFile.copy()
    ratios[ratios.columns[rel_cols:]] =
myFile[myFile.columns[rel_cols:]].div(myFile[ref_col] + 1, axis = 0)
    log2 = myFile.copy()
    log2[log2.columns[rel_cols:]] =
ratios[ratios.columns[rel_cols:]].apply(np.log2)
    return log2
```

#### 2.3 median-smoothing functions

```
def medsmooth(x,y,rel_cols):
    return x[x.columns[rel_cols:]].apply(signal.medfilt,args=(y,))

def smoothing(log2, chrcol = "chr", rel_cols = 7):
    smoothres1, smoothres2, smoothres3 = log2.copy(), log2.copy(),
log2.copy()
    smoothres1.iloc[:,rel_cols:] = log2.groupby(by =
chrcol).apply(medsmooth,11,rel_cols)
    smoothres2.iloc[:,rel_cols:] = smoothres1.groupby(by =
chrcol).apply(medsmooth,21,rel_cols)
    smoothres3.iloc[:,rel_cols:] = smoothres2.groupby(by =
```

```
chrcol).apply(medsmooth,41,rel_cols)
    return smoothres1, smoothres2, smoothres3
```

#### 2.4 post-smoothing per-sample normalization

```
def Xfilt(log2, smooth, chrcol = "chr", rel_cols = 7):
    Xchr = log2[log2[chrcol]==23]
    Xchr = Xchr[Xchr.columns[rel_cols:]]
    Xmedians = Xchr.apply(np.median)
    Xadj = myFile.copy()
    Xadj[Xadj.columns[rel_cols:]] =
smooth[smooth.columns[rel_cols:]].subtract(Xmedians)
    return Xadj

def nonXfilt(log2, smooth, chrcol = "chr", rel_cols = 7):
    nonXchr = log2[log2[chrcol]<23] # can peel off higher chrs with
frequent CN
    nonXchr = log2[log2[chrcol]>0]  # can peel off lower chrs with
frequent CN
    nonXchr = nonXchr[nonXchr.columns[rel_cols:]]
    nonXmedians = nonXchr.apply(np.median)
    nonXadj = myFile.copy()
    nonXadj[nonXadj.columns[rel_cols:]] =
smooth[smooth.columns[rel_cols:]].subtract(nonXmedians)
    print(nonXmedians)
    return nonXadj
```

#### 2.5 cytoband average per-gene CN estimates

```
def make_cbmeans(adjusted, cband, rel_cols,
               rcn_genes_fn="./esca_tcga/data_linear_CNA.txt",
               rcn_gene_col='Hugo_Symbol',
               ecn_gene_col='ID', chr_col='chr', inCHRloc =
"inCHRloc"):
    #rcn_genes = set(pd.read_csv(rcn_genes_fn, sep="\t").dropna()
[rcn_gene_col])
    #ecn_genes = set(Xadj[ecn_gene_col])
    #common_genes = rcn_genes & ecn_genes
    #Xadj_c = Xadj[[x in common_genes for x in Xadj[ecn_gene_col]]]
    #Xadj_c.reset_index(drop = True, inplace = True)
    adjusted.insert(rel_cols+1,"cband_start",np.NaN)
    ecn_cb = adjusted.groupby(by = chr_col).apply(assign_cytoband,
cband, chr_col, inCHRloc).dropna()
    #e_cbmeans = ecn_cb.iloc[:,rel_cols:].groupby(by =
"cband_start").mean()
    e_cbmeans = ecn_cb.iloc[:,rel_cols:].groupby(by = 'cband_start',
sort = False).mean() # to order by chr, then by position
    return e_cbmeans
```

##### 2.6 convert cytoband CN estimates to ordinal gain\loss\2n categories

```
# to collapse cytoband averaged CN to ~ relative copy number
# thresholds (thresh) are for STRONG GAIN, weak gain and weak loss,
STRONG LOSS
# usual values would be 0.4 and -0.4 or 0.5 and -0.5 for strong
changes
# as well as 0.2 to 0.25 and -0.2 to -0.25 for the weak thresholds
# GISTIC uses 0.2 and 0.4 levels for these threholds, be we find
CoFrEE to be more sensitive and so prefer 0.25 and 0.5 as thresholds
# the weak threshold (change vs no change) is more important than the
strong (some vs stronger change)
# no change cytobands (~2n) will be those between the weak thresholds,
e.g < 0.25 and > -0.25
def grade_cn(e_cbmeans):
    def val2cat(x, thresh=None):
        if thresh is None:
            thresh=[-0.5,-0.225,0.225,0.5]
            levels = [-2,-1,0,1,2]
        for thr,lev in zip(thresh,levels):
            if x<thr: return lev
        return levels[-1]
    return e_cbmeans.applymap(val2cat)
```

##### 2.7 graphing functions

```
def heatmap(inpt, title = "estimated cb_means", vmin = -2, vmax = 2,
cmap='RdYiBu_r', figsize=(3,5), **kw):
    fig = plt.figure(figsize=figsize, dpi=150)
    r = sns.heatmap(inpt, vmin = vmin, vmax = vmax, center = 0,
cmap=cmap, **kw)
    r.set_title(title)
    return fig
```

#### 3. Various Test Functions

##### 3.1 get working directory path

```
import os #provides operating system dependent functionality
currentDirectory = os.getcwd()
print(currentDirectory)
```

```
/Users/derekn/Documents/work 2021/CN from expn/Code discussion/May
2021 code
```

##### 3.2 show cytoband (cband) table

```
cbfile = "cytoBand.txt"
# just to test cband import & adjust to big bands
```

```
cband = getbands(cbfile)
cband
```

```
    cbname   chr       start        end  chrnum
0     1p36  chr1           0   27600000       1
1     1p35  chr1    27600000   34300000       1
2     1p34  chr1    34300000   46300000       1
3     1p33  chr1    46300000   50200000       1
4     1p32  chr1    50200000   60800000       1
..     ...   ...         ...        ...     ...
315   Xq27  chrX   138900000  148000000      23
316   Xq28  chrX   148000000  156040895      23
317   Yp11  chrY           0   10400000      24
318   Yq11  chrY    10400000   26600000      24
319   Yq12  chrY    26600000   57227415      24

[320 rows x 5 columns]
```

#### 3.3 load and view sample data file

```
#### INPUT FILE CHARACTERISTICS ####
```

Cohort datafile should have:

- unique gene ID
- be ordered by chromosome and chromosomal position we do this by ranking chr * 1,000,000,000 + start
- gene symbol
- chromosome band location
- chr (as a number plus 'X' and 'Y', or 23 and 24 - for human)
- start location (base)
- end location (base)
- at least one reference data column
  - these samples must be from the same expression study as tumors to per-gene expression levels compare
  - average of several non-cancer samples from the same tissue as tumors
  - average of cancer samples without site-specific changes, or fewest changes [known copy number data needed]
- sample data files (which can included individual reference samples)

```
#### ABOUT THE EXAMPLE COHORT ####
```

For the below example, we use a refined publicly released meningioma cohort to compare with CASPER.

The data is presented in the Yale maningioma cohort, GEO series Series https://www.ncbi.nlm.nih.gov/geo/query/acc.cgi?acc=GSE85133 published in Clark et al., 2016:https://pubmed.ncbi.nlm.nih.gov/27548314/.

For simplicity we manually reduced samples to those presented in CaSpER paper by Harmanci et al., in Fig 3A https://www.nature.com/articles/s41467-019-13779-x: tumors with chromosome 22 (NF2) deletions. We manually extracted gene expression data presented in the CaSpER R-library, rather than GEO to ensure direct comparison.

This cohort of RNAseq data are provided as part of the "yale_meningioma.rda" within CaSpER R library (https://github.com/akdess/CaSpER/blob/master/demo/meningioma.R).

We chose this cohort as an example because it provides a fair comparion between the more simplistic approach of CoFrEE (eg no BAF component) against the well accepted CaSpER approach. The advantage CoFrEE offers is that it works on both array and RNAseq cohorts.

It is for these reasons we chose this cohort as the contrast to CaSpER in Fig 1B and Sup Fig 4. Data in our paper shows that CoFrEE echos the findings of CaSpER for chr 22q deletion, but is overly sensitive in specific samples in other parts of the genome.

```
# gse = GEOparse.get_GEO(filepath="./GEO_data/GSE85133_family.soft",
silent=False)

# myFile = pd.read_csv('meningioma cohort from CaSpER.csv') # all
samples
myFile = pd.read_csv('meningioma sample with chr22 deletion v2.csv') #
sample subset in CASPER pre-accepted version figure
myFile
```

```
       ensembl gene ID        myLOC    symbol chromosome   chr
start  \
0      ENSG00000187634   1000925738    SAMD11    1p36.33     1
925737
1      ENSG00000188976   1000944204     NOC2L    1p36.33     1
944203
2      ENSG00000187961   1000960587    KLHL17    1p36.33     1
960586
3      ENSG00000187583   1000966497   PLEKHN1    1p36.33     1
966496
4      ENSG00000187642   1000975204     PERM1    1p36.33     1
975203
...                ...          ...       ...        ...   ...         ..
.
15804  ENSG00000008735  22050600685  MAPK8IP2   22q13.33    22
50600684
15805  ENSG00000100299  22050625018      ARSA   22q13.33    22
50625017
15806  ENSG00000251322  22050674642    SHANK3   22q13.33    22
50674641
15807  ENSG00000100312  22050738196       ACR   22q13.33    22
50738195
15808  ENSG00000079974  22050767501    RABL2B   22q13.33    22
50767500

            end  NOchange5mean  NOchange5median    MN-1171  MN-
```

```
60835  \
0         944581         9.821590          9.613405  10.708870  9.909009

1         959290         9.299376          9.050384   9.211174  9.041787

2         965715         5.186108          5.138979   6.317308  6.068920

3         975108         2.884740          2.639051   5.209350  4.646615

4         982093         1.498832          1.604236   3.321832  3.860656

...          ...              ...               ...        ...       ...

15804  50611551         8.133005          8.064420   6.909788  5.897643

15805  50628173         8.358584          8.750986   8.845384  7.723959

15806  50733210         8.061165          7.335456   6.812072  7.606485

15807  50745334         1.192346          1.032696   1.700366  1.323653

15808  50783663         7.623634          7.443624   7.943874  7.994384


        MN-1236   MN-1237   MN-1137   MN-1161     MN-60      MN-5
0      9.449096  9.135786  8.618603  8.893933  8.006841  8.602705
1      9.451082  9.563796  9.645113  8.817251  8.525805  9.085696
2      5.438287  5.094681  4.885604  5.852335  5.108745  5.769310
3      1.548657  3.019341  2.413422  1.243690  1.570899  3.132956
4      1.959092  2.791462  1.846561  1.243690  0.000000  2.410869
...         ...       ...       ...       ...       ...       ...
15804  7.552810  7.361798  7.606338  8.712935  6.693302  7.839126
15805  8.649231  8.095251  8.226837  7.010235  8.061081  8.471966
15806  7.011699  6.047809  5.628380  7.757993  8.183497  8.083764
15807  0.000000  1.127410  0.899541  0.751907  0.000000  1.447409
15808  6.212705  6.599283  6.600916  6.646487  7.805496  6.812537

[15809 rows x 17 columns]
```

### 4. MAIN FUNCTION

```
''' main analysis function has the following assumptions
      - loci are in chromosome and chromosomal order
      - expression values are non-log format
      - CN is based on aneuploidy, as compared to base genome (2n)
      - by-cytoband assigned a single CN for a band, with no thought
for partial or multiple CN involvment
        this is to provide an overall pattern; by-chromosome view
provides the by-locus CN
```

```
    and has the following steps:
      - for each locus in each sample generate a log2 ratio, compared
to chosen reference
        (usually mean/median of normal tissue samples)
      - 3 rounds of median smoothing (filtering) are done, reportable
in smoothmed1, 2 & 3
      - either mean of X chromosome or mean of all autosomal ratios
are used to adjust CN for each sample
          this is essential to account for experimental and ploidy
level differences.
      - generates a by-cytoband CN value for each sample (mean CN of
genes that start in a given cytoband)
'''

def cofree_main(myFile, ref_col="NOchange5mean", rel_cols = 10,
                chr_col = "chr", cbfile = "./cytoBand.txt",
                inCHRloc = "inCHRloc",
                rcn_genes_fn="./esca_tcga/data_linear_CNA.txt",
                rcn_gene_col='Hugo_Symbol',
                ecn_gene_col='ID',cmap = 'Spectral',vmin = -3, vmax =
3,
                figure_fn_template=None):
    log2 = logFC(myFile, ref_col, rel_cols)
    smooth3 = smoothing(log2, chr_col, rel_cols)[2] #there are 3
median smoothed versions in this dataframe ([0] smoothres1 to [2]
smoothres3)
#    Xadj = Xfilt(log2, smooth3, chr_col, rel_cols)
    nonXadj = nonXfilt(log2, smooth3, chr_col, rel_cols)

    cband = getbands(cbfile)
    e_cbmeans =
make_cbmeans(smooth3,cband,rel_cols,rcn_genes_fn,rcn_gene_col,ecn_gene
_col,chr_col, inCHRloc)
#    e_cbmeans =
make_cbmeans(Xadj,cband,rel_cols,rcn_genes_fn,rcn_gene_col,ecn_gene_co
l,chr_col, inCHRloc)
#    e_cbmeans =
make_cbmeans(nonXadj,cband,rel_cols,rcn_genes_fn,rcn_gene_col,ecn_gene
_col,chr_col, inCHRloc)

    cbm_fig = heatmap(e_cbmeans, "estimated cytoband means",
cmap=cmap, vmin=vmin, vmax=vmax)
    e_cbmeans_graded = grade_cn(e_cbmeans) # thresh=[-0.5,-
0.225,0.225,0.5]
    cbmg_fig = heatmap(e_cbmeans_graded, "graded copy numbers per
cytoband", cmap=cmap, vmin=-2, vmax=2)
    e_cbmeans.to_csv('bigband_medians.txt', sep='\t', index=False)
    return {'cn':nonXadj, 'cb_means':e_cbmeans,
'cb_grades':e_cbmeans_graded,
            'cb_means_fig':cbm_fig, 'cb_grades_fig':cbmg_fig}
```

```
res = cofree_main(myFile, ref_col="NOchange5median", rel_cols=9,
chr_col="chr", inCHRloc="start",
                        vmin=-0.25, vmax=0.25, cmap="RdYlBu_r")

MN-1171     -0.182794
MN-60835    -0.195442
MN-1236     -0.140653
MN-1237     -0.153379
MN-1137     -0.147455
MN-1161     -0.151463
MN-60       -0.174347
MN-5        -0.169872
dtype: float64
```

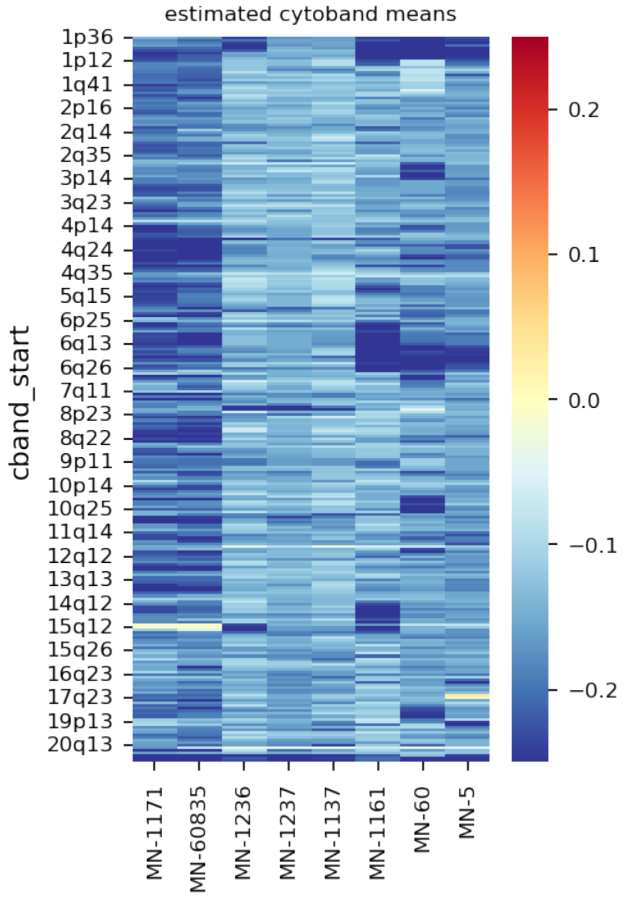

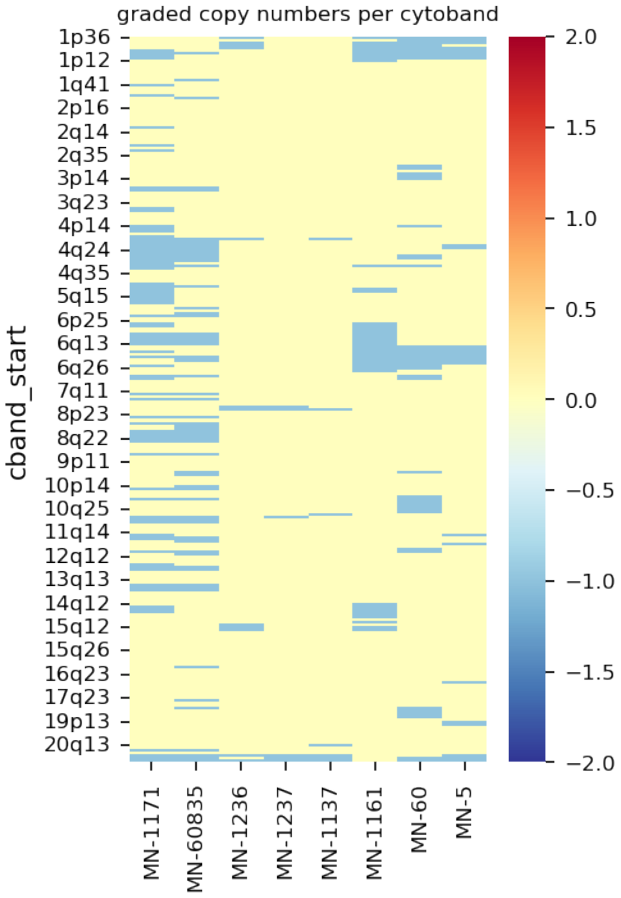


### 5. Output File Manipulations

simple code lines to copy outputs, make subsets and output to a text file

#### 5.1 isolate output tables

```
cb_grades = res["cb_grades"].copy() # copy number grades for each
cytoband (bigband)

cb_means = res["cb_means"].copy() # av. cn value for each cytoband
(bigband)
```

#### 5.2 isolate bands with changes

```
#cytoband subset with at least one sample with a non-zero grading
graded_non_zero = cb_grades.loc[cb_grades.index[(cb_grades !=
0).any(axis=1)],:]

#cytoband subset with at least one sample with a non-zero mean
non_zero = cb_means.loc[cb_means.index[(cb_means.abs() >
0.05).any(axis=1)],:]
```

#### 5.3 copy key table to file

```
# copy cytoband mean values to a file
cb_means.to_csv('bigband_averages.txt', sep='\t', index=True)
cb_grades.to_csv('bigband_cn_grades.txt', sep='\t', index=True)
```

#### 5.4 isolate specific regions

```
# cytoband means for chr22q only
cb1 = cb_means.loc[cb_means.index.str.startswith("22q")] # just chr
22q
cb1
```

```
              MN-1171  MN-60835   MN-1236   MN-1237   MN-1137   MN-
1161  \
cband_start

22q11       -0.273393 -0.348250 -0.293175 -0.292703 -0.296315 -
0.221942
22q12       -0.267438 -0.354858 -0.221261 -0.256681 -0.241386 -
0.201984
22q13       -0.255843 -0.278796 -0.231982 -0.284129 -0.287489 -
0.208293

               MN-60      MN-5
cband_start
22q11       -0.128403 -0.239348
22q12       -0.252538 -0.277701
22q13       -0.264221 -0.269671
```

```
# cytoband means for chr22q only
cb2 = cb_grades.loc[cb_grades.index.str.startswith(('20', '21',
'22'))] # just chr 20-22
cb2
```

```
             MN-1171  MN-60835  MN-1236  MN-1237  MN-1137  MN-1161
MN-60  \
cband_start

20p13              0         0        0        0        0        0
0
20p12              0         0        0        0        0        0
0
20p11              0         0        0        0        0        0
0
20q11              0         0        0        0        0        0
0
20q12              0         0        0        0        0        0
0
20q13              0         0        0        0       -1        0
0
```

```
21q11                0         0        0        0        0        0
0
21q21               -1        -1        0        0        0        0
0
21q22                0         0        0        0        0        0
0
22q11               -1        -1       -1       -1       -1        0
0
22q12               -1        -1        0       -1       -1        0
-1
22q13               -1        -1       -1       -1       -1        0
-1

             MN-5
cband_start
20p13           0
20p12           0
20p11           0
20q11           0
20q12           0
20q13           0
21q11           0
21q21           0
21q22           0
22q11          -1
22q12          -1
22q13          -1
```

```
# show cn grades for the first and last 5 bands - python defaults
cb_grades
```

```
             MN-1171  MN-60835  MN-1236  MN-1237  MN-1137  MN-1161
MN-60  \
cband_start

1p36                0         0       -1        0        0       -1
-1
1p35                0         0        0        0        0        0
-1
1p34                0         0       -1        0        0       -1
-1
1p33                0         0       -1        0        0       -1
-1
1p32                0         0       -1        0        0       -1
-1
...               ...       ...      ...      ...      ...      ...
...
21q21              -1        -1        0        0        0        0
0
21q22               0         0        0        0        0        0
```

```
0
22q11                  -1        -1        -1        -1        -1         0
0
22q12                  -1        -1         0        -1        -1         0
-1
22q13                  -1        -1        -1        -1        -1         0
-1

             MN-5
cband_start
1p36           -1
1p35           -1
1p34           -1
1p33            0
1p32           -1
...           ...
21q21           0
21q22           0
22q11          -1
22q12          -1
22q13          -1

[277 rows x 8 columns]
```

### 6.0 Heatmeap Graphic Configurations

code lines to modify or output heatmap figures

#### 6.1 graded non-zero band heatmap

```
#cytoband subset with at least one sample with a non-zero grading
graded_non_zero = cb_grades.loc[cb_grades.index[(cb_grades !=
0).any(axis=1)],:]

fig = heatmap(graded_non_zero, cmap = "Spectral", linewidths = 0,
linecolor = (.5,.5,.5), xticklabels = True, vmin = -2, vmax = 1)
```

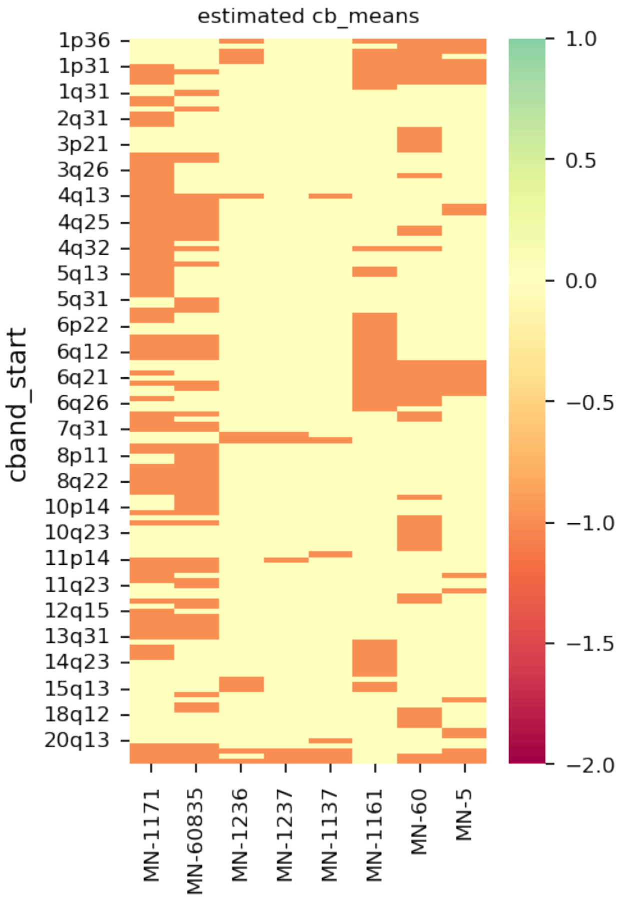


```
fig.savefig("graded_nonzero_bands.png",  bbox_inches = "tight")

!cd
```

#### 6.2 plot two chr heatmap per graph

```
# provides regular expression matching operations similar to those
found in Perl
import re

#just bands for chr19 to chr22 - two chrs per graph
chroms = [(f"meningioma_cbmeans_{c}-{c+1}", f"({c}|{c+1})[pq]") for c
in range(19,22,2)]
chroms

[('meningioma_cbmeans_19-20', '(19|20)[pq]'),
 ('meningioma_cbmeans_21-22', '(21|22)[pq]')]

for base,rx in chroms:
    fig = heatmap(cb_means[cb_means.index.str.match(rx)],
                         cmap="Spectral",
                         linewidths=0.1, linecolor=(0.5,0.5,0.5),
```

```
                              yticklabels=True, vmax = 0.5, vmin = -0.5
                             )
    fig.savefig(base+".png", bbox_inches = "tight")
```

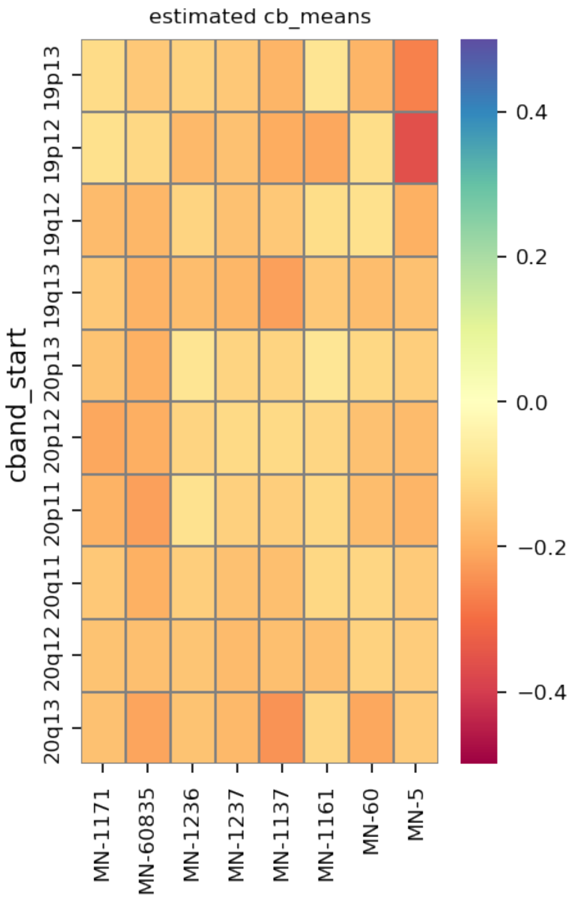

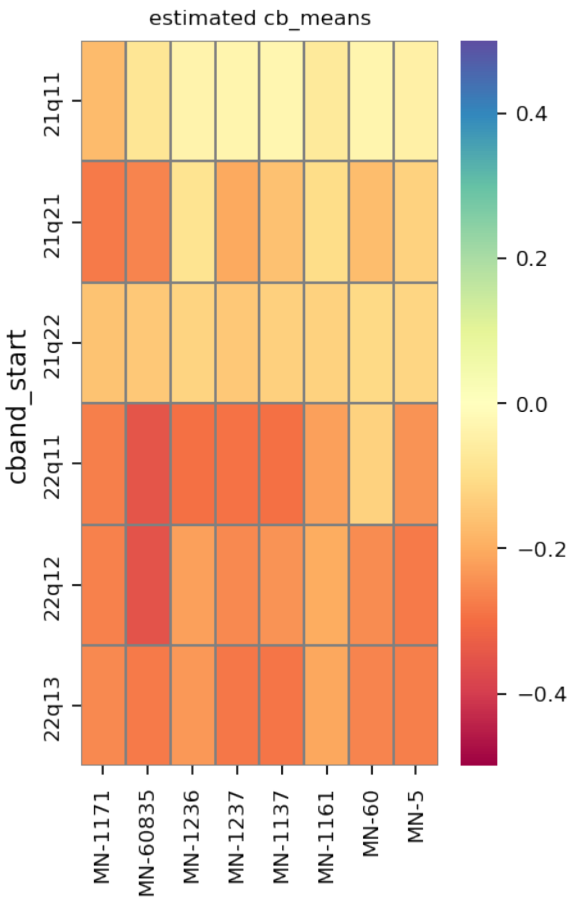


```
cb_grades

             MN-1171  MN-60835  MN-1236  MN-1237  MN-1137  MN-1161
MN-60  \
cband_start

1p36               0         0       -1        0        0       -1
-1
1p35               0         0        0        0        0        0
-1
1p34               0         0       -1        0        0       -1
-1
1p33               0         0       -1        0        0       -1
-1
1p32               0         0       -1        0        0       -1
-1
...              ...       ...      ...      ...      ...      ...
...
21q21             -1        -1        0        0        0        0
0
21q22              0         0        0        0        0        0
0
```

```
22q11              -1        -1       -1       -1       -1        0
0
22q12              -1        -1        0       -1       -1        0
-1
22q13              -1        -1       -1       -1       -1        0
-1

             MN-5
cband_start
1p36           -1
1p35           -1
1p34           -1
1p33            0
1p32           -1
...           ...
21q21           0
21q22           0
22q11          -1
22q12          -1
22q13          -1

[277 rows x 8 columns]
```
